# DeepBIO is an automated and interpretable deep-learning platform for biological sequence prediction, functional annotation, and visualization analysis

**DOI:** 10.1101/2022.09.29.509859

**Authors:** Ruheng Wang, Yi Jiang, Junru Jin, Chenglin Yin, Haoqing Yu, Fengsheng Wang, Jiuxin Feng, Ran Su, Kenta Nakai, Quan Zou, Leyi Wei

## Abstract

Here, we present DeepBIO, the first-of-its-kind automated and interpretable deep-learning platform for high-throughput biological sequence functional analysis. DeepBIO is a one-stop-shop web service that enables researchers to develop new deep-learning architectures to answer any biological question. Specifically, given any biological sequence data, DeepBIO supports a total of 42 state-of-the-art deep-learning algorithms for model training, comparison, optimization, and evaluation in a fully automated pipeline. DeepBIO provides a comprehensive result visualization analysis for predictive models covering several aspects, such as model interpretability, feature analysis, functional sequential region discovery, *etc*. Additionally, DeepBIO supports 9 base-level functional annotation tasks using deep-learning architectures, with comprehensive interpretations and graphical visualizations to validate the reliability of annotated sites. Empowered by high-performance computers, DeepBIO allows ultra-fast prediction with up to million-scale sequence data in a few hours, demonstrating its usability in real application scenarios. Case study results show that DeepBIO provides an accurate, robust, and interpretable prediction, demonstrating the power of deep learning in biological sequence functional analysis. Overall, we expect DeepBIO to ensure the reproducibility of deep-learning biological sequence analysis, lessen the programming and hardware burden for biologists, and provide meaningful functional insights at both sequence-level and base-level from biological sequences alone. DeepBIO is publicly available at http://inner.wei-group.net/DeepBIO.

## Introduction

The development of next-generation sequencing techniques has led to an exponential increase in the amount of biological sequence data accessible, including genomic, transcriptomic, and proteomic sequences, *etc*. It naturally poses an important challenge - how to build the relationships from sequences to structures and functions (1). Given such large-scale data, traditional wet-lab experimental methods are laborious, time-consuming, and high-cost for functional analysis. Alternatively, the booming development of machine learning approaches has paved a new way to understand the complex mapping from biological sequences to their structures and functional mechanisms. Over the past few decades, data-driven machine learning approaches have emerged as powerful methods to enable the automated and fast function prediction ab initio from sequences alone, providing a new perspective on studying the functional aspects of biological sequences (2–5).

The increasing use of machine-learning workflows in biological sequence analysis and the willingness to disseminate trained machine learning models have pushed computer scientists to design more user-friendly solutions. In recent years, there is an increasing number of web servers and software packages developed for this purpose. For instance, BioSeq-Analysis is the first platform to analyze various biological sequences via machine learning approaches (6). Later, Liu et al. established BioSeq-Analysis2.0 (7) to automatically generate various predictors for biological sequence analysis at both residue level and sequence level. Additionally, iLearnplus is a popular platform that provides a pipeline for biological sequence analysis, which comprises feature extraction and selection, model construction, and analysis of prediction results (8). More recently, Liu et al. developed BioSeq-BLM, a web platform that introduces different biological language models for DNA, RNA and protein sequence analysis (9). Besides, other representative tools include iFeatureOmega (10), Rcpi (11), and protr (12), *etc*. The platforms and tools have boosted the use of machine learning solutions for biological sequence analysis tasks in biology. However, these traditional machine-learning workflows have some drawbacks. For instance, to train a good model, strong professional knowledge is usually required in terms of designing feature descriptors, selecting machine learning algorithms, as well as setting up the model parameters, which limit their usability in real applications to some extent; on the other hand, they cannot support large-scale prediction and analysis. Recently, deep learning has played a complementary role over traditional machine learning due to its excellent scalability and adaptivity. Therefore, some deep-learning analysis tools have been developed, such as Kipoi (13), Pysster (14), and Selene (15). Kipoi (13) is a repository of ready-to-use trained models for genomic data analysis. Pysster (14) is a Python package that supports training deep-learning models only with convolutional neural networks on biological sequence data. More recently, Chen et al. developed Selene, a PyTorch-based deep-learning library for quick and simple development, training, and application of deep-learning model architectures for biological sequence data (15).

These tools make the use of deep learning more convenient for data-driven biological sequence analysis to some extent. However, there remain some key challenges that need to be addressed. First, despite the availability of deep-learning tools, they are not completely automated. Running the tools still faces some technical challenges for non-expert researchers. On one hand, it requires strong professional knowledge and programming skills to set up sophisticated software, which directly limits the generic use of deep learning in nonexpert communities (i.e., researchers without any computer science training). On the other hand, it is extremely difficult for researchers to train deep-learning models with the enormous scale of data space, since the major downside of deep learning is its computational intensity, requiring high-performance computational resources and long training time. Accordingly, a web platform that enables an automated deep-learning pipeline is highly required. Second, most of the existing tools cannot meet the high demand of the research community, since they provide few deep-learning architectures (seen in **Supplementary Table 1**), such as convolutional neural networks, for model construction. In fact, there are some state-of-the-art deep-learning models with successful applications in bioinformatics problems. For example, Graph Neural Networks demonstrated their ability to address complex biological problems, for example, the prediction of microRNA-disease interaction (16). Large-scale language pre-trained models like DNABERT and ProtBERT, showed strong capacities in discriminative feature learning in biological sequence analysis (5,17). Thus, it is necessary to enable the use of state-of-the-art deep-learning approaches in biology. Ultimately, existing tools lack comprehensive result analysis, which might limit users’ understanding of what the deep-learning models learn; how reliable the deep-learning predictions are; and why the deep learning performs well.

In this work, we present DeepBIO, an automated deep-learning platform for biological sequence prediction, functional annotation, and result visualization analysis. To be specific, our DeepBIO distinguishes itself from other platforms with the following unique advantages: **(1) One-stop-shop service.** DeepBIO is the first-of-its-kind platform as a code-free web portal to ensure the reproducibility of deep-learning biological sequence analysis and to lessen the programming burden for biologists. **(2) Pure deep-learning platform.** DeepBIO is a purely deep-learning platform that integrates over 40 state-of-the-art mainstream deep-learning algorithms, including convolutional neural networks, advanced natural language processing models, and graph neural networks (detailed in **Supplementary Table 2**), which enables researchers of interest to train, compare, and evaluate different architectures on any biological sequence data. **(3) Two predictive modules.** DeepBIO is the first platform that supports not only sequence-level function prediction for any biological sequence data, but also allows 9 base-level functional annotation tasks using pre-trained deep-learning architectures, covering DNA methylation, RNA methylation, and protein-binding specificity. Besides, we further offer an in-depth comparative analysis between the predictions by our models and by experimental data to validate the reliability of the predictions. Notably, empowered by high-performance computers, we demonstrate that DeepBIO supports fast prediction with up to million-scale sequence data, demonstrating its usability in real application scenarios. **(4) Comprehensive result visualization analysis**. Aiming to help researchers to dig into more insights from the model predictions, DeepBIO offers a comprehensive visualization result analysis with a variety of interactive figures and tables, covering the following five aspects: statistics analysis of input data; prediction models’ performance evaluation and comparison; feature importance and interpretable analysis; model parameter analysis; and sequence conservation analysis. In particular, integrating interpretable mechanisms (*e.g*., attention heatmap, and motif discovery) into deep-learning frameworks enables researchers to analyze which sequential regions are important for the predictions, addressing the issue of the “black box” in deep learning and effectively building the relationship between biological sequences to functions.

## Materials and Methods

### The overall framework of DeepBIO

The DeepBIO platform fully automates the model training process and applies more than forty deep-learning approaches for sequence classification prediction and functional site analysis based on genomic and proteomic data. **Figure 1A** illustrates the overall framework of the proposed DeepBIO, which consists of four modules: (1) Data input module, (2) Sequence-level functional prediction module, (3) Base-level functional annotation module, and (4) Result report module. The workflow of DeepBIO is described as follows. Firstly, DeepBIO takes biological sequence data from the data input module, where we can handle three biological sequence types including DNA, RNA, and protein. Next, there are two functional modules for two types of tasks respectively; one is the sequence-level functional prediction module for binary classification tasks, and the other is the base-level functional annotation module for functional annotation tasks. The detailed web techniques and the overview of the interaction between the front end and back end are shown in **Figure 1B**. In the sequence-level functional prediction module, we support users in automatically training, evaluating, and comparing the deep-learning models with their input data. Specifically, there are four main steps (seen in **Figure 1C**) in this module: (I) Data pre-processing, (II) Model construction, (III) Model evaluation, and (IV) Visualization analysis. In the functional annotation module, we provide the base-level functional annotation using deep-learning approaches, such as DNA methylation, RNA methylation, and protein-binding specificity prediction. Similarly, we also curated four steps for this module (seen in **Figure 1C**): (I) Data selection, (II) Task selection, (III) Model loading, and (IV) Result visualization. Ultimately, in Result report module, we provide a series of visualizatioin analysis results with various kinds of data formats. Below, we emphasize the details of the four modules.

**Figure 1.**
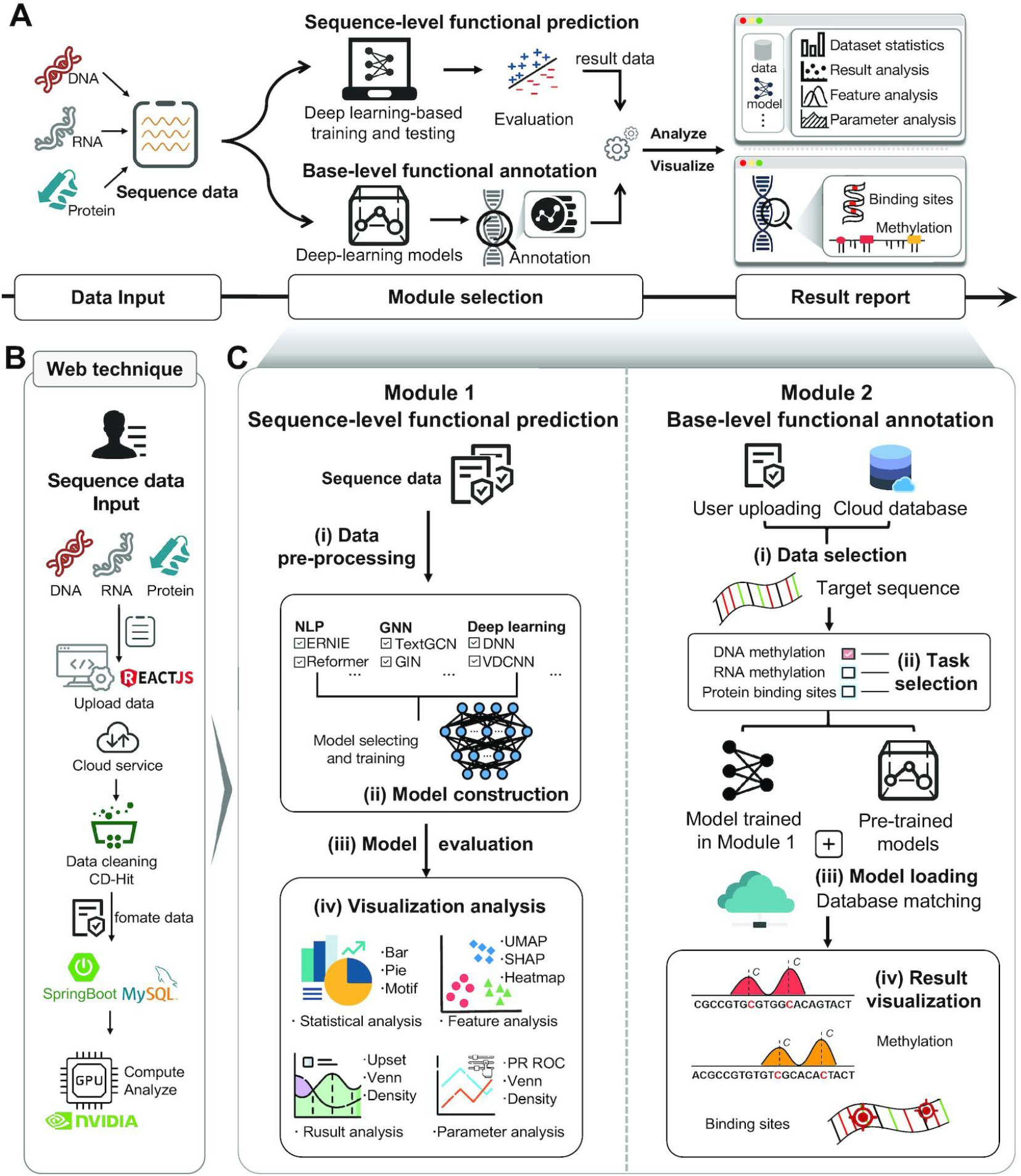
The organization of the DeepBIO platform. **(A)** The architecture of the DeepBIO platform. DeepBIO contains four modules, including the data input module, the sequence-level functional prediction module, the base-level functional annotation module, and the result report module. DeepBIO accepts DNA, RNA, and protein sequence data from the data input module, with some additional user-selectable settings (*e.g*., sequence similarity setting, imbalanced data processing, and data augmentation). The sequence-level functional prediction module provides the deep-learning biological sequence binary classification service while the base-level functional annotation module supports the prediction of functional sites for user-provided sequences. In the result report module, DeepBIO provides user-friendly, interactive, and interpretable visualization representations to deliver results of the above two functional modules as a download-free report. **(B)** The web technique of DeepBIO and an overview of the framework. **(C)** The workflow of two main functional modules. The sequence-level functional prediction module consists of four steps: (I) Data pre-processing, (II) Model construction, (III) Model evaluation, and (IV) Visualization analysis. DeepBIO outputs classification predictions of input sequences based on the four steps and displays the result reports with interactive graphs and tables; The base-level functional annotation module includes the following steps: (I) Data selection, (II) Task selection, (III) Model loading, and (IV) Result visualization. DeepBIO offers multiple methylation annotations for DNAand RNA sequences while supporting ligand-binding site recognition for protein sequences.

### Data input module

DeepBIO serves as an automatically integrated analysis platform based on biological sequence data, including DNA, RNA, and protein, which can be entered in the online input box or uploaded with the standard file format. Besides, users can select deep-learning models and different settings (*e.g*., whether to turn on the data enhancement option). After obtaining the user’s input data, DeepBIO further performs data cleaning to ensure that the user input data is ready and legal to pass into the next module and run properly.

### Sequence-level functional prediction module

#### Step 1. Data pre-processing

This step is an optimization-based data processing phase that uses a variety of approaches to handle and obtain the input sequence data that is suitable for model training, ensuring the robustness of the final results and improving the success rate of our service. We design four sections to pre-process the input data: (I) Sequence legitimacy testing, (II) Sequence similarity setting, (III) Imbalanced data processing, and (IV) Data augmentation. In Section (I), for the input sequence data, we first examine whether the input data has the legal data format (*e.g*., FASTA), and then check if the input sequences contain illegal characters (*e.g*., other characters except for A, T, C, G in DNA sequences). In Section (II), considering that the high sequence similarity in the input data might lead to the performance evaluation bias, we thus provide a commonly used tool named CD-HIT (18) to reduce the sequence similarity in the input data. The similarity threshold ranges from 0.7 to 1. In Section (III), considering that a majority of datasets in real scenarios might exist the data imbalance issue (i.e., the number of the positives and negatives are extremely imbalanced), we further provide several imbalanced data processing strategies that are commonly used in the machine learning field, including Focal loss (19), Synthetic Minority Oversampling Technique (SMOTE) (20), and Adaptive Synthetic (ADASYN) (21). By doing so, we can prevent the biased prediction of models due to the imbalanced data distribution and perhaps achieve the improvement in the overall predictive performance. Finally, in Section (IV), to deal with the few sample learning problem, we further provide some data augmentation methods, such as sequence replacement, sequence flipping, and sequence cropping, etc, which can improve the robustness and generalization ability of the deep-learning models.

#### Step 2. Model construction

The pre-processed sequence data is then fed into user-selected models to train and make the binary prediction. For the scenario that users do not provide sequence data with training and testing labels, we randomly split the dataset into the training and testing set by a ratio of 8:2 for model training and evaluation. **Table 1** summarizes all the state-of-the-art deep-learning models in our platform. There are a total of 42 deep-learning models, covering classic deep-learning methods (*e.g*., Deep Neural Networks (DNN), and Gate Recurrent Unit (GRU), *etc*), Natural Language Processing (NLP) methods (*e.g*., Transformer, and Bidirectional Encoder Representations from Transformers (BERT), etc.), and Graph Neural Network (GNN) methods (*e.g*., Graph Convolutional Network (GCN)). Notably, each of the above models can be exploited to deal with all three types (DNA, RNA, and protein) of sequence analysis tasks. In addition, DeepBIO permits users to select multiple models, train, and compare their performance on the same datasets. It’s worth noting that we have also deployed several deep-learning models (*e.g*., DNABERT, RNABERT, and ProtBERT) that are well pre-trained on large-scale biological background sequences. They can be utilized for fine-tuning the models on small datasets, if necessary, to alleviate the overfitting problem and increase the models’ generalization capability.

**Table 1.**
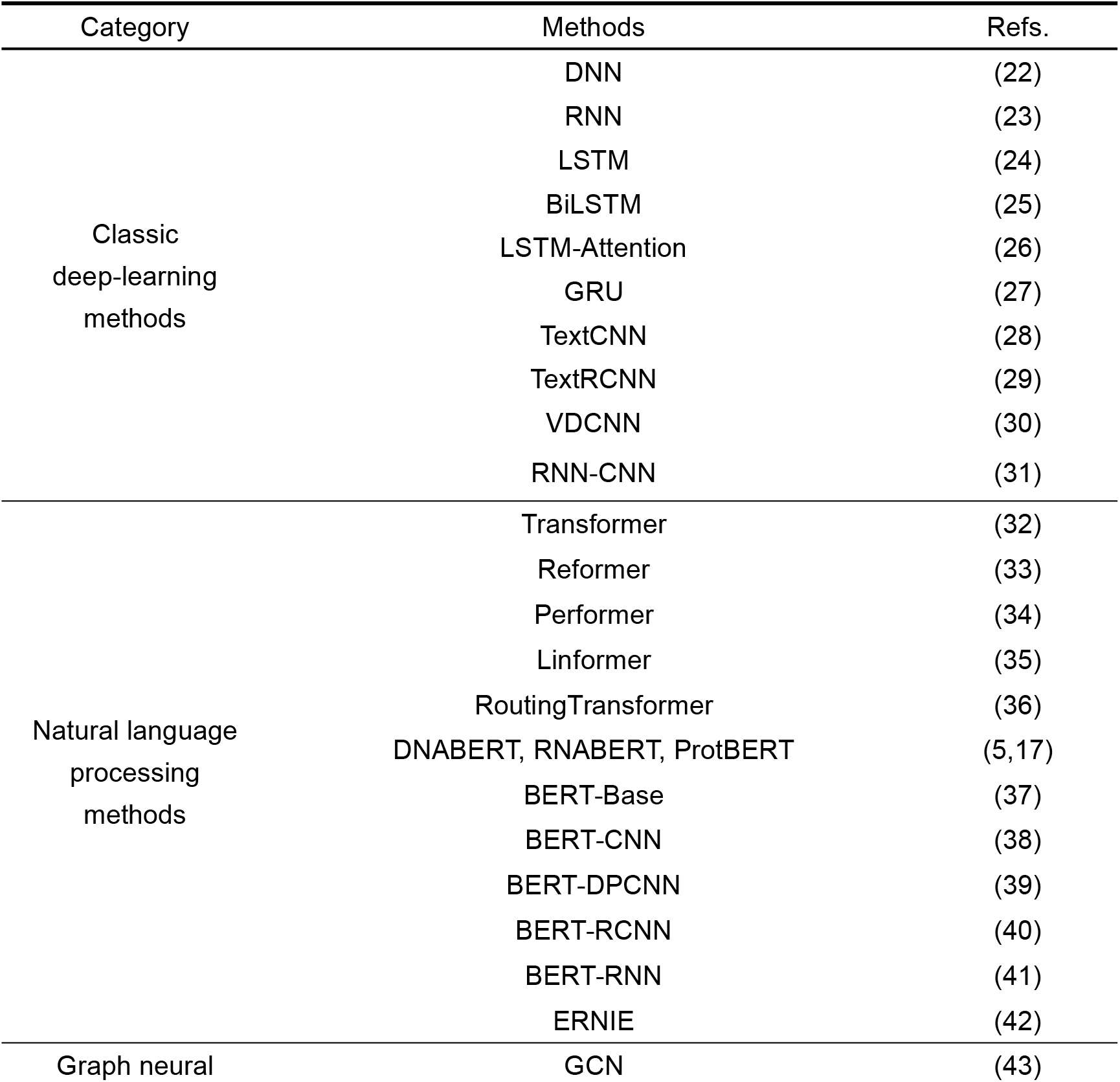
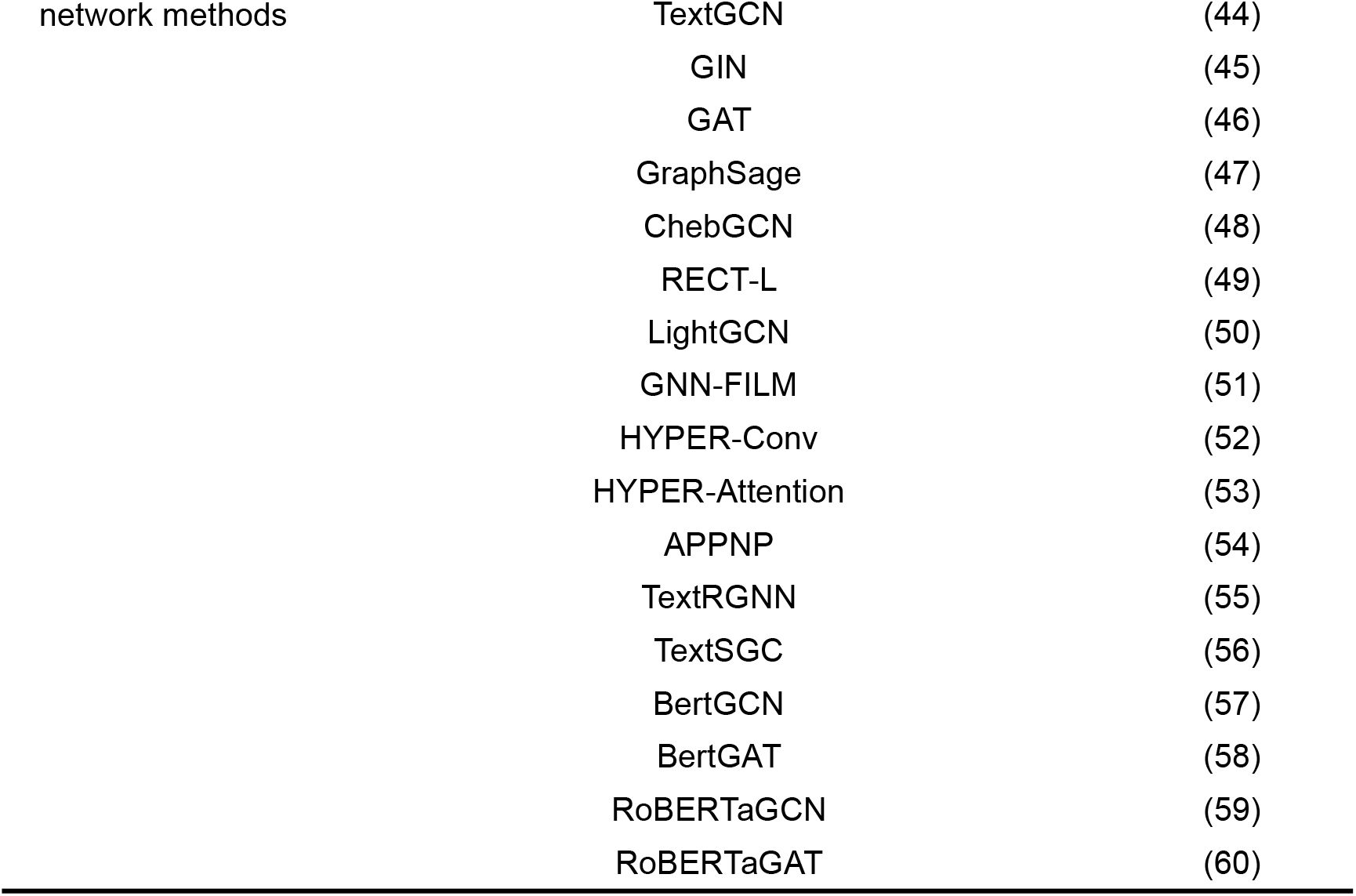
Deep-learning models in DeepBIO.

#### Step 3. Model evaluation

For the performance comparison of deep-learning models, we choose several commonly used evaluation metrics, including Sensitivity (SN), Specificity (SP), Accuracy (ACC), and Matthews’ Correlation Coefficient (MCC). The formulas of these metrics are described as follows:

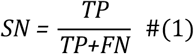

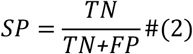

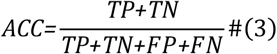

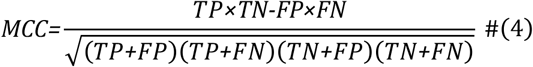

where TP, FP, TN, and FN represent the numbers of true positives, false positives, true negatives, and false negatives, respectively. For the comprehensive performance comparison of different models, the Area Under Receiver Operating Characteristic curve (AUC) and Area Under Precision Recall Curve (AP), which range from 0 to 1, are calculated based on the ROC curve and the PR curve, respectively. The ROC curve shows the proportion of true positives versus false positives, in which the AUC equals the probability of ranking a randomly chosen true target higher than a randomly chosen decoy target. The PR curve is used more frequently to evaluate the performance of a model on imbalanced datasets (61). Similarly, we can calculate the AP, which equals the average of the interpolated precisions. The higher the AUC and AP values, the better the predictive performance of the underlying model.

#### Step 4. Visualization analysis

DeepBIO combines the prediction results and intermediate data analysis generated from above, and helps users to better understand the input data through the statistical analysis and the model results through various types of visualization analyses. To provide users with intuitive and comprehensive result analysis, we design and illustrate multiple visual presentation formats, including the pie plot, histogram, sequence logo graph, ROC and PR curves, kernel density plot, and scatter plot. In **Table 2**, we present the four sections of visualization analysis, which are dataset statistical analysis, result analysis, feature analysis, and parameter optimization analysis. The details of each section are described as follows.

**Table 2.**
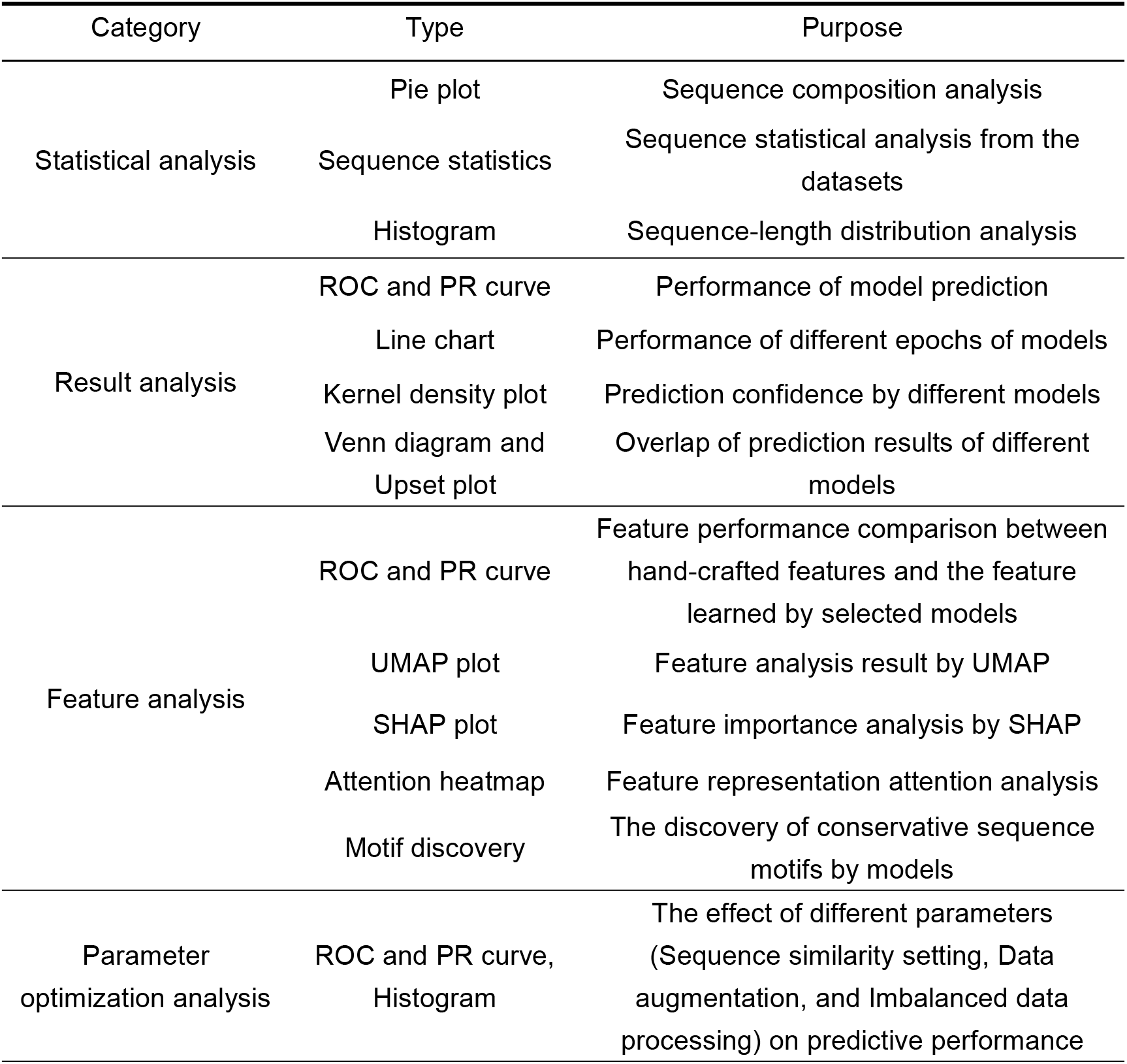
Result visualization analysis in the prediction module.

For dataset statistical analysis, we summarize the overall status of the input datasets to assist users in better comprehending the datasets themselves. There are the following three statistical analyses, including sequence composition analysis, sequence motif analysis, and the sequence-length distribution analysis. In sequence composition analysis, DeepBIO plots the sequence composition histogram for the samples in the positive and negative dataset. In sequence motif analysis, users can obtain the conservation of each position along the sequences. Ultimately, the sequence-length distribution analysis provides users with the length preference of the sequences in the dataset.

For result analysis, the DeepBIO visualization analysis includes plots considering all the evaluation metrics presented in step 3, to conduct detailed performance comparisons among different models, such as ROC and PR curves, the performance of different epochs, density distribution of model prediction confidence, *etc*. Besides, we offer a Venn diagram and an Upset plot to illustrate the relationship among different models from the overlapping prediction results. It can be concluded that in this part users can get an intuitive comparison of prediction performance for different deep-learning models.

For feature analysis, we use a dimensionality reduction tool called Uniform Manifold Approximation and Projection (UMAP) (62) for feature space analysis, and a commonly used tool Shapley Additive explanation (SHAP) (63) for feature importance analysis. UMAP can map and visualize the high-dimensional features learned from the deep-learning models to the low-dimensional space. It intuitively illustrates how good the learned features are. On the other hand, SHAP gives a good explanation of the feature importance, which quantitively measures the impact of each of the learned features on the predictive performance. Besides, for some specific models like DNABERT and RNABERT, to further explore what the model learns, we employ the attention heatmap to visualize the information the model captures from either global or local perspectives. Furthermore, the motif discovery enables users to know the conservative sequential determinants the models learn from the datasets. The details of the motif discovery process are described as follows. Firstly, we apply the attention mechanism of BERT-based models to identify key regions from the sequences. Then, for those identified regions, we further extract and visualize the corresponding motifs using the attention scores obtained from the model. In this way, we highlight the concerns of different models for different features and enhance the interpretability of the deep-learning models.

For parameter optimization analysis, we provide users with some parameter options to make their own choice to optimize the model training, including the effect of different sequence similarities in datasets, the effect of different data augmentation strategies, and the effect of different imbalanced data processing methods on predictive performance. Besides, we also give some tabular information, including a summary of the input datasets and the performance of deep-learning models. This helps the users to compare the results of the same model with different parameters.

### Base-level functional annotation module

The base-level functional annotation module is to predict the functions of the biological sequences at base level using deep learning. In this module, we provide 9 annotation tasks, including the methylation annotation for DNA and RNA sequences, and the ligand-binding site recognition for protein sequences. The module comprises four steps: Data selection, Task selection, Model loading, and Result visualization. The details of this module are as follows.

#### Step 1. Data selection

In this step, DeepBIO provides users to upload the sequences they want to annotate. Particularly, for the DNA methylation site prediction task, we support two ways to input sequence data, that is (1) uploading sequence data by themselves, (2) and selecting a segment of our preset human genome by choosing a specific cell line and chromosome.

#### Step 2. Task selection

There are three types of functional annotation tasks: DNA methylation site prediction, RNA methylation site prediction, and protein-ligand binding site prediction. For DNA methylation site prediction, there are DNA N^4^-methylcytosine (4mC) site prediction, DNA 5-hydroxymethylcytosine (5hmC) site prediction, and DNA N^6^-methyladenine (6mA) site prediction. For RNA methylation site prediction, there are RNA 5-hydroxymethylcytosine (5hmC) site prediction, RNA N^6^-methyladenosine (m6A) site prediction, and RNA N^4^-methylcytidine (m4C) site prediction. For protein-ligand binding site prediction, there are DNA-binding site prediction, RNA-binding site prediction, and Peptide-binding site prediction.

#### Step3. Model loading

In this step, there are two modes for deep-learning model selection. One is default and the other is customized. For the default mode, we provide several deep-learning models that are well pre-trained with large-scale biological data, such as DNABERT. The well-pre-trained models generally have good generalization ability and achieve good performance for different downstream tasks. For the customized mode, we allow users to upload the models they trained for the same tasks in the sequence-level functional prediction module.

#### Step 4: Result visualization

In this part, we provide users with the in-depth annotation result analysis based on the model predictions and biological experiment data (*e.g*., histone modification signals and protein 3D structural information). Specifically, for DNA, we designed two visualization sections, including functional site annotation and integrative analysis. For functional site annotation, we predict the functional sites that match the selected task type (*e.g*., DNA 4mC site prediction) and perform the model-predicted confidence of the corresponding position. In the integrative analysis, we perform statistical analysis between the expression of DNA methylation and different histone modification signals. For RNA, due to the constraint of data, we only perform the functional site annotation section same as in DNA. In addition, for protein, to intuitively show the binding site annotation results, we visualize and annotate the binding residues on the 3D structure of a given protein sequence with PDB ID.

### Result report module

We list the analysis results in the form of reports, which involve various kinds of data formats from the results of the two main functional modules. The users can view or download freely all of the result data and charts. Notably, each submitted task is immediately notified to the corresponding user once it is completed.

## Results and discussion

### Case Study

We showcase real-world applications of DeepBIO with two bioinformatic scenarios: (1) the prediction of DNA 6mA methylation and (2) the prediction of the protein toxicity. We emphasize that the underlying objective is to illustrate how to use our platform for two such diverse applications, rather than securing the top predictive performance compared to the state-of-the-art methods.

DNA methylation is closely associated with a variety of key processes, ranging from genomic imprinting, and aging, to carcinogenesis (64). Therefore, for the task of the DNA 6mA methylation prediction, we use the *D.melanogaster* dataset with 11,191 methylation sequences (as positives) and 11,191 non-methylation sequences (as negatives) as an example dataset (65) for DNA sequence functional analysis. In addition, protein toxicity is closely related to most neurodegenerative diseases and accurate toxicity prediction leads to significant promotion in new drug design. We choose a protein toxicity benchmark dataset (66), select protein sequences between 50 and 1200 amino acids, and obtain the final example dataset with 3,379 toxic animal protein sequences (as positives) and 5,464 non-toxic animal protein sequences (as negatives) for protein functional analysis. With the two example datasets, we first randomly split them into the training set and testing set and then input them into our online platform to demonstrate the data analysis and various functions of DeepBIO. More details of the visualization results of the two cases are described below.

### Dataset statistics

**Figure 2** illustrates the dataset statistics of the protein toxicity dataset. The proportion and number of bases or amino acids in the dataset are counted and illustrated as a pie plot and composition histogram, respectively (**Figure 2A**), from which we can clearly see the difference between the positives and the negatives from the compositional perspective. In addition, we calculate and show the sequence length distribution (**Figure 2B**), which intuitively gives users the sequence-length preference in the datasets. For example, it can be seen from **Figure 2B** that positive samples are mainly distributed in the range of 50 to 150 amino acids, while the negative samples have a relatively even distribution. Furthermore, we use a motif analysis tool called Weblogo to generate the sequence motifs of the datasets, which enables users to analyze the composition and conservation at the sequence level (**Figure 2C**).

**Figure 2.**
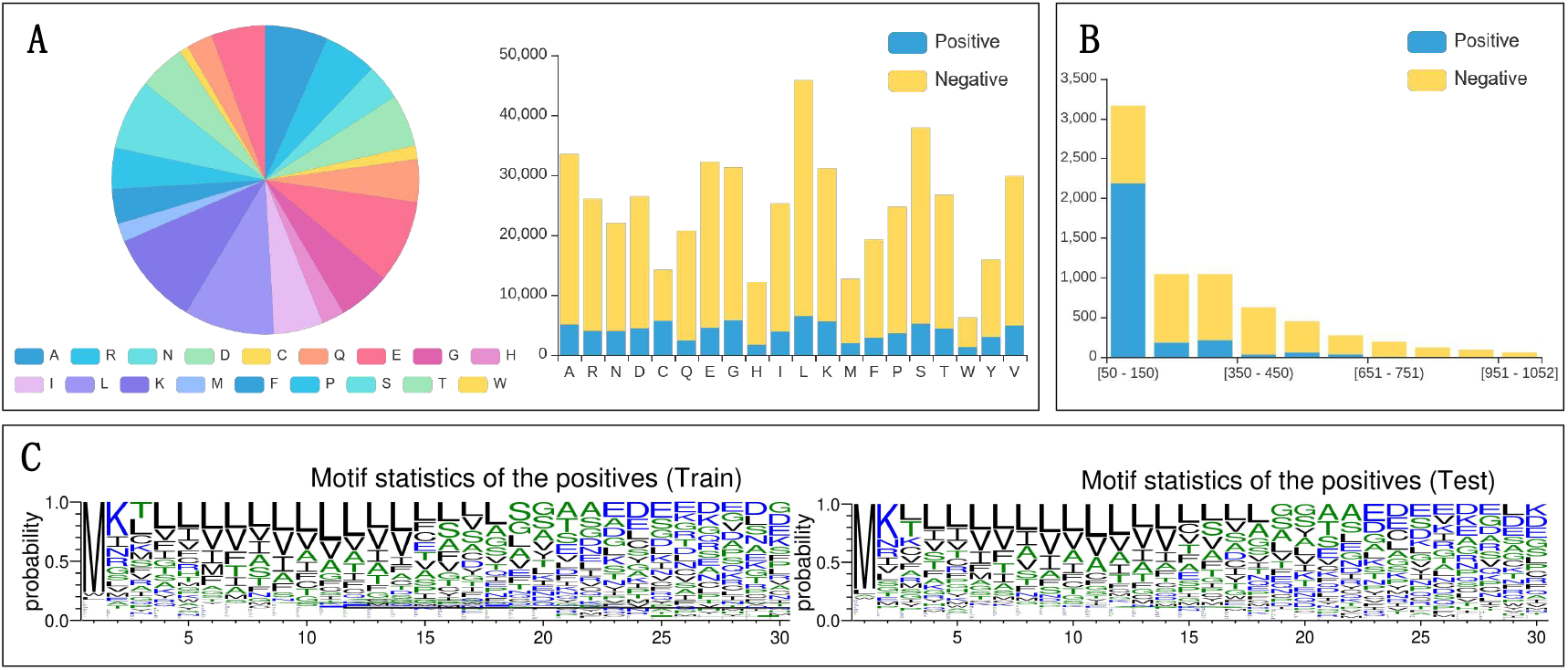
The statistical analysis on the protein toxicity dataset. **(A)** Sequence compositions of positive and negative samples in the training and testing sets; **(B)** Sequence length distribution of positive and negative samples in the training and testing sets; **(C)** Motifs of sequence statistics from the training and testing sets.

### Model prediction analysis

DeepBIO provides detailed analysis for users to interpret and show comprehensive performance comparisons among different deep-learning models. Here, we select four models (DNABERT, DNN, LSTM, and RNN) to train and compare their prediction performance on the DNA methylation dataset. **Figure 3A** shows the performances of the compared models in terms of ACC, Sensitivity, Specificity, AUC, and MCC. It can be seen that DNABERT achieves the best performance with the ACC, Sensitivity, Specificity, AUC, and MCC of 0.921, 0.896, 0.946, 0.967, and 0.844, respectively. For a better evaluation of models, **Figure 3B** illustrates the ROC and PR curves of the compared models. As seen in **Figure 3B**, DNABERT achieved the highest AUC and AP values of 0.968 and 0.973, respectively, which further confirms that DNABERT is better than the other compared models. To further study the generalization ability of the compared models, we draw the epoch plot in **Figure 3C** to show the trends of accuracy and calculation loss for each model during the training process. From **Figure 3C**, it is easy to find that except for DNN, the accuracy of the other three models (DNABERT, LSTM, and RNN) increases rapidly at first and then the accuracy curves get smooth gradually. Meanwhile, their calculation losses converge to lower values, indicating that these models are well trained and their predictions are reliable. It is worth noting that during the training process, DNN always achieves relatively low accuracy, however with high calculation loss on the example dataset (seen in **Figure 3C**). This indicates that it is not a good and robust model after training. In addition, to intuitively show the relationship among different models from their prediction results, we further provide an Upset plot and a Venn diagram in **Figure 3D-E**. From **Figure 3D-E**, we can see that most positive samples are successfully predicted from the overlapping predictions of the compared models. Moreover, to clearly understand the prediction preference for each model, we visualize the density distribution of the prediction confidence of different deep-learning models in **Figure 3F**. As can be seen, DNN mainly focuses its prediction confidence on around 0.5, which is the center part of the *X*-axis, illustrating its poor discrimination ability between positive and negative samples on the example dataset. On the contrary, the other three models (DNABERT, LSTM, and RNN) achieve higher prediction confidence both on positive and negative samples, indicating that they possess better classification ability on the example dataset. The comparison results for this part on the dataset of protein toxicity can be seen in **Supplementary Figures 1-6.**

**Figure 3.**
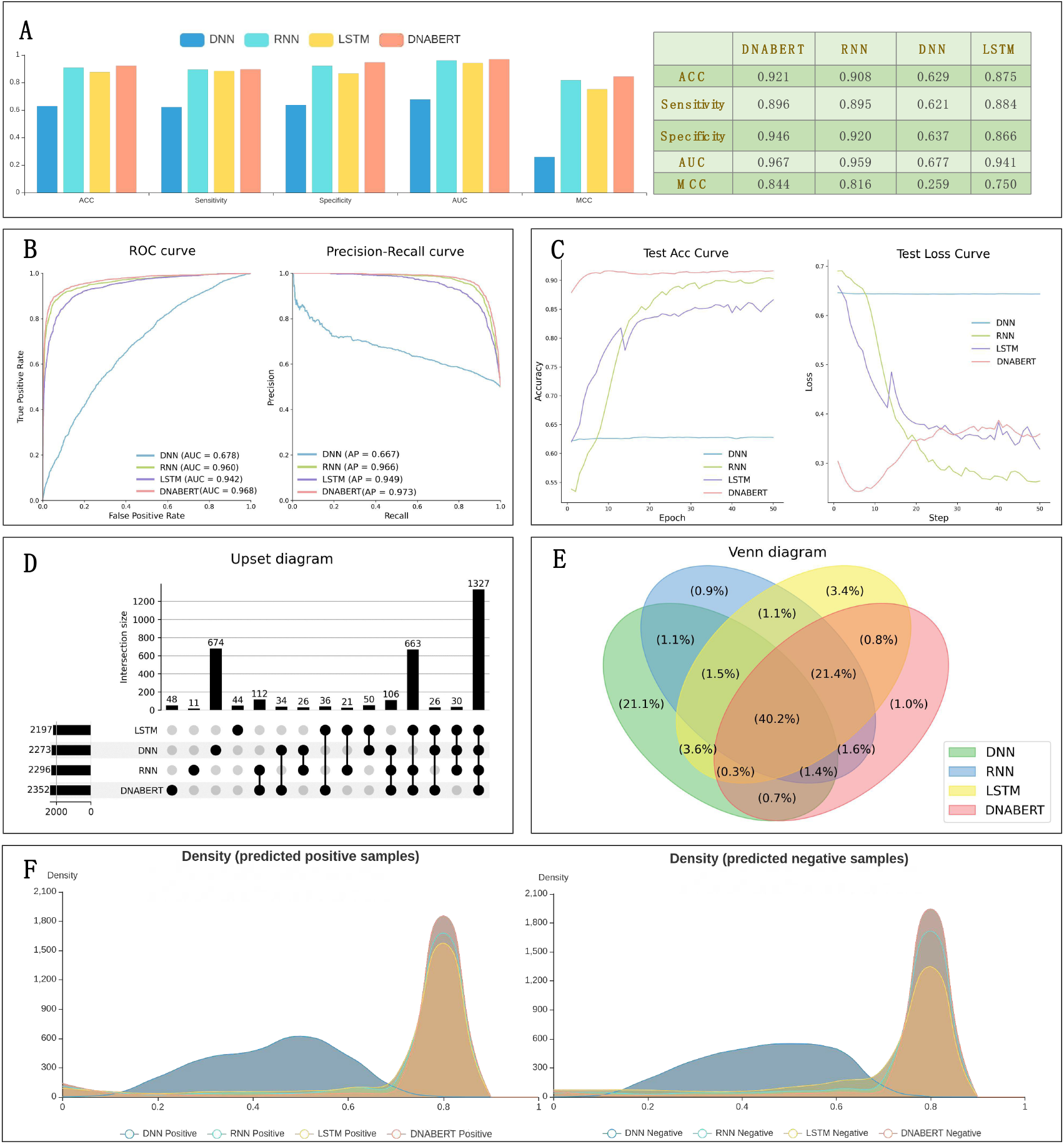
The prediction result analysis from different models using DeepBIO. **(A)** Performance comparison among different models in terms of ACC, Sensitivity, Specificity, AUC, and MCC; **(B)** ROC and PR curves of different deep-learning models; **(C)** The trends of performance and calculation loss with each epoch, showing ACC and loss change process on different models; **(D-E)** Upset plot and Venn diagram to express the relationships of prediction results for different models; **(F)** Density distribution of the prediction confidence for different deep-learning models.

### Feature analysis and visualization

To compare the features automatically learned by deep-learning models with the traditional hand-crafted features, we select some commonly used feature encodings, including Accumulated Nucleotide Frequency (ANF) (67), binary (68), Composition of K-spaced Nucleic Acid Pairs (CKSNAP) (69), and Dinucleotide Composition (DNC) (70,71) for nucleotide sequences, and show the prediction performance in terms of ACC, Sensitivity, Specificity, AUC, MCC, and AP in **Figure 4A-B**. It is worth noting that in **Figure 4A-B**, the compared deep-learning models except for DNN outperformed most of the traditional feature encodings, allowing us to easily conclude that the features learnt from the deep-learning methods generally are superior to hand-crafted features.

**Figure 4.**
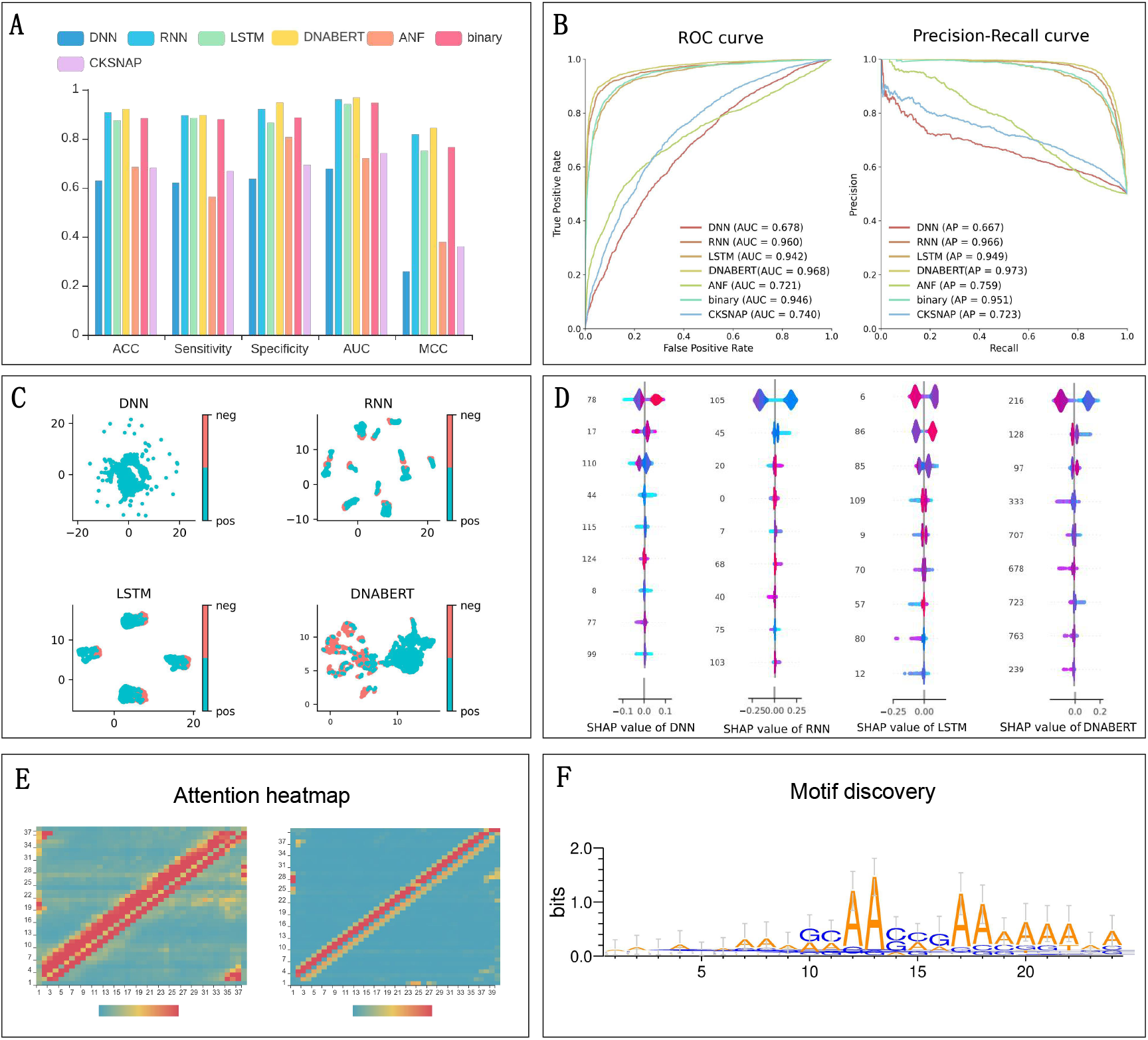
The visualization of feature analysis and model interpretation using DeepBIO. **(A)** Prediction performance comparison between deep-learning models and traditional hand-crafted feature-based methods; **(B)** ROC and PR curves of deep-learning models and traditional hand-crafted feature-based methods; **(C)** UMAP feature visualization results of different deep-learning models; **(D)** SHAP feature importance analysis of different deep-learning models; **(E)** Attention heatmap of the user-choosing biological sequence; **(F)** Motif discovery by deep-learning models.

In addition, to intuitively evaluate the feature representations learnt from different deep-learning models, we employ UMAP to conduct the dimension reduction and feature visualization (**Figure 4C**). In **Figure 4C,** each point represents a sample in the dataset, and the positive and negative classes are distinguished by different colors. In this case, we can get a better sense of the classification ability of each model from the UMAP plot. Specifically, **Figure 4C** shows that compared with the other three models (DNABERT, LSTM, and RNN), the samples belonging to different classes in the feature space of DNN are almost connected, making it difficult to distinguish the positives from the negatives. Moreover, the samples of different classes are more clearly distributed in the feature space of DNABERT than in the feature space of RNN and LSTM. In addition, we also provide the SHAP plot (**Figure 4D**), trying to assign each feature a value of importance for the model prediction. Each row represents the SHAP value distributions of a feature, and the *X*-axis refers to the SHAP value, where the value of SHAP >0 shows that the prediction favors the positive class, and a value <0 indicates that the prediction tends to be of the negative class. The color of sample points in the figure indicates the corresponding feature value; redder points mean higher feature values, while bluer points indicate lower feature values. The features are sorted according to the sum of SHAP values incorporating all the samples in the dataset. Therefore, the SHAP plot shows the relationship between the value of features and their impact on the model prediction. From **Figure 4D** we can see that the high-dimension features play a predominant role for DNABERT and the top-ranked feature values in DNABERT are more easily distinguished in the color compared with other models, indicating that DNABERT performs better in feature representation learning.

To improve the interpretability of the deep-learning models, in **Figure 4E**, we calculate the importance of the base at each position on the user-given sequence from the attention matrix in the model and visualize it with normalized attention scores, enabling us to analyze which region the model considers more important for the prediction. To further study what the models learn from the training process, we extract the important regions with high attention scores through the attention mechanism of the models to find conservative sequence motifs in **Figure 4F** from the datasets, which might be related to biological functions. The detailed feature analysis results on the dataset of protein toxicity can be seen in **Supplementary Figures 7-9**.

### Functional annotation analysis

To intuitively illustrate in-depth annotation result analysis of biological sequences, we provide users with functional site prediction and result visualization analysis as in **Figure 5**. As seen in **Figure 5A**, we can predict DNA methylation sites with corresponding positions and prediction confidences based on user-provided DNA sequences or a segment of our preset human genome. In addition, if users select a segment from our preset human genome by choosing a specific cell line and chromosome, we will display the statistical analysis between our DNA methylation site predictions and relevant histone modification signals queried from the database (**Figure 5B**), assisting in mining the potential relations between them. Moreover, for protein sequence annotation tasks, we predict and visualize the binding site annotation results on the 3D structure of the given protein sequence with its PDB ID. Specifically, **Figure 5C** shows the protein-peptide binding site prediction results of an example protein (PDB ID 1a81A), and the residues in red color on the protein 3D structure indicate the protein-peptide binding sites.

**Figure 5.**
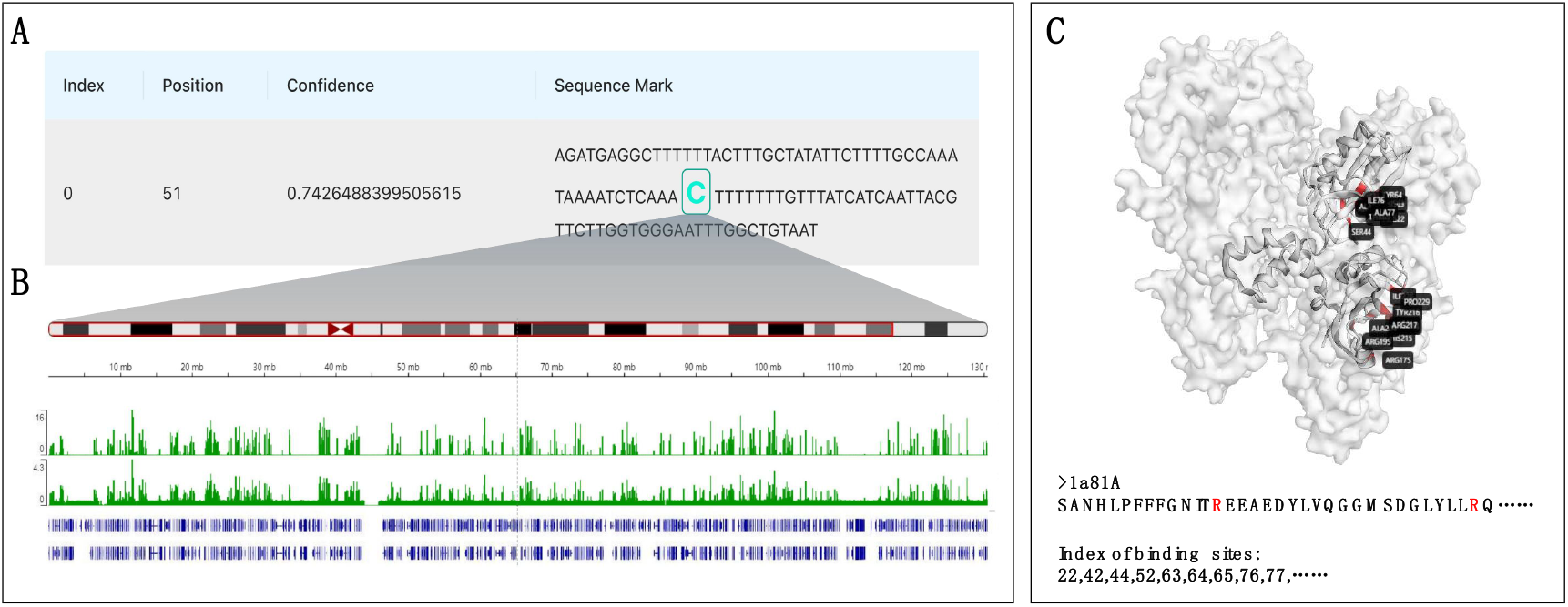
The biological sequence functional annotation analysis using DeepBIO. **(A)** Prediction results of DNA methylation site annotation tasks; **(B)** Comparison and analysis between the DNA methylation site prediction results and relevant histone modification signals; **(C)** Prediction results of protein-binding site annotation tasks.

### Webserver Implementation

DeepBIO is empowered by the professional high-performance graphic processing architecture with million-scale calculating capability. Specifically, DeepBIO is deployed with Ubuntu 18.04 Linux system, multiple Intel Xeon Silver 4210R CPUs, 256GB RAM, hundreds of TB Solid State Drive, and NVIDIA RTX 3090 GPU clusters. Unlike other online platforms that use CPU for training models and making predictions, the models we provide are all trained and evaluated based on GPU, which significantly reduces the computational time. It can be seen from **Supplementary Table 3** that we can spend less time optimizing deep-learning models and enable fast and accurate predictions. In addition, **Supplementary Figures 10-15** show that we have carefully designed the front-end page to give users an interactive visualization of the results while they can download each result graph freely. To be specific, we use React framework to set up the user interface, Spring Boot framework for server implementation in the back end, python and R programming languages for the model’s construction and visualization, and MySQL database to manage the data storage. Besides, to make a better Web experience for users, we also constructed interactive graphs to visualize the analysis result on the front page. Our platform can run stably on many browsers including Internet Explorer (≥v.7.0), Mozilla Firefox, Microsoft Edge, Safari, and Google Chrome. In conclusion, with our DeepBIO platform, data scientists and researchers limited by equipment resources can now employ high-performance computers to tackle their most challenging work.

## Availability of Data and Materials

As an online platform for biological sequence analysis, DeepBIO can be freely accessed without registration via http://inner.wei-group.net/DeepBIO. All code used in data analysis and preparation of the manuscript, alongside a description of necessary steps for reproducing results, can be found in a GitHub repository accompanying this manuscript: https://github.com/WLYLab/DeepBIO. The source code for DeepBIO is also freely available at Zenodo website (DOI: 10.5281/zenodo.7547847).

## Author contributions

L.W. conceived the basic idea and designed the DeepBIO framework. R.W., Y.J, J.J., C.Y., H. Y., F.W., and J.F. set up the server and visualized the result analysis. Y.J., R.W., L.W., and C.Y. designed the interface and tested the server. R.W., C.Y., J.J., Q.Z., and L.W. wrote the manuscript. K.N., R.S., Q.Z., and L.W. revised the manuscript.

## Reference

1. Larranaga, P., Calvo, B., Santana, R., Bielza, C., Galdiano, J., Inza, I., Lozano, J.A., Armananzas, R., Santafe, G., Perez, A. et al. (2006) Machine learning in bioinformatics. Brief Bioinform, 7, 86–112.

2. Wang, R., Jin, J., Zou, Q., Nakai, K. and Wei, L. (2022) Predicting protein–peptide binding residues via interpretable deep learning. Bioinformatics, 38, 3351–3360.

3. Jiang, Y., Wang, R., Feng, J., Jin, J., Liang, S., Li, Z., Yu, Y., Ma, A., Su, R., Zou, Q. et al. (2022) Explainable deep graph learning accurately modeling the peptide secondary structure prediction. bioRxiv doi: https://www.biorxiv.org/content/10.1101/2022.06.09.495580v2, 10 August 2022, preprint: not peer reviewed.

4. Jin, J., Yu, Y., Wang, R., Zeng, X., Pang, C., Jiang, Y., Li, Z., Dai, Y., Su, R. and Zou, Q. (2022) iDNA-ABF: multi-scale deep biological language learning model for the interpretable prediction of DNA methylations. Genome biology, 23, 1–23.

5. Elnaggar, A., Heinzinger, M., Dallago, C., Rehawi, G., Wang, Y., Jones, L., Gibbs, T., Feher, T., Angerer, C., Steinegger, M. et al. (2021) ProtTrans: Towards Cracking the Language of Lifes Code Through Self-Supervised Deep Learning and High Performance Computing. IEEE Transactions on Pattern Analysis and Machine Intelligence, 1–1.

6. Liu, B. (2019) BioSeq-Analysis: a platform for DNA, RNA and protein sequence analysis based on machine learning approaches. Briefings in bioinformatics, 20, 1280–1294.

7. Liu, B., Gao, X. and Zhang, H. (2019) BioSeq-Analysis2. 0: an updated platform for analyzing DNA, RNA and protein sequences at sequence level and residue level based on machine learning approaches. Nucleic acids research, 47, e127–e127.

8. Chen, Z., Zhao, P., Li, C., Li, F., Xiang, D., Chen, Y.Z., Akutsu, T., Daly, R.J., Webb, G.I., Zhao, Q. et al. (2021) iLearnPlus: a comprehensive and automated machine-learning platform for nucleic acid and protein sequence analysis, prediction and visualization. Nucleic Acids Res, 49, e60.

9. Li, H.-L., Pang, Y.-H. and Liu, B. (2021) BioSeq-BLM: a platform for analyzing DNA, RNA and protein sequences based on biological language models. Nucleic acids research, 49, e129–e129.

10. Chen, Z., Liu, X., Zhao, P., Li, C., Wang, Y., Li, F., Akutsu, T., Bain, C., Gasser, R.B., Li, J. et al. (2022) iFeatureOmega: an integrative platform for engineering, visualization and analysis of features from molecular sequences, structural and ligand data sets. Nucleic Acids Research, 50, W434–W447.

11. Cao, D.-S., Xiao, N., Xu, Q.-S. and Chen, A.F. (2015) Rcpi: R/Bioconductor package to generate various descriptors of proteins, compounds and their interactions. Bioinformatics, 31, 279–281.

12. Xiao, N., Cao, D.-S., Zhu, M.-F. and Xu, Q.-S. (2015) protr/ProtrWeb: R package and web server for generating various numerical representation schemes of protein sequences. Bioinformatics, 31, 1857–1859.

13. Avsec, Ž., Kreuzhuber, R., Israeli, J., Xu, N., Cheng, J., Shrikumar, A., Banerjee, A., Kim, D.S., Beier, T. and Urban, L. (2019) The Kipoi repository accelerates community exchange and reuse of predictive models for genomics. Nature biotechnology, 37, 592–600.

14. Budach, S. and Marsico, A. (2018) Pysster: classification of biological sequences by learning sequence and structure motifs with convolutional neural networks. Bioinformatics, 34, 3035–3037.

15. Chen, K.M., Cofer, E.M., Zhou, J. and Troyanskaya, O.G. (2019) Selene: a PyTorch-based deep learning library for sequence data. Nature methods, 16, 315–318.

16. Li, C., Liu, H., Hu, Q., Que, J. and Yao, J. (2019) A novel computational model for predicting microRNA–disease associations based on heterogeneous graph convolutional networks. Cells, 8, 977.

17. Ji, Y., Zhou, Z., Liu, H. and Davuluri, R.V. (2021) DNABERT: pre-trained Bidirectional Encoder Representations from Transformers model for DNA-language in genome. Bioinformatics.

18. Li, W. and Godzik, A. (2006) Cd-hit: a fast program for clustering and comparing large sets of protein or nucleotide sequences. Bioinformatics, 22, 1658–1659.

19. Lin, T.-Y., Goyal, P., Girshick, R., He, K. and Dollár, P. (2017), Proceedings of the IEEE international conference on computer vision, pp. 2980–2988.

20. Chawla, N.V., Bowyer, K.W., Hall, L.O. and Kegelmeyer, W.P. (2002) SMOTE: synthetic minority over-sampling technique. Journal of artificial intelligence research, 16, 321–357.

21. He, H., Bai, Y., Garcia, E.A. and Li, S. (2008), 2008 IEEE international joint conference on neural networks (IEEE world congress on computational intelligence). IEEE, pp. 1322–1328.

22. LeCun, Y., Bengio, Y. and Hinton, G. (2015) Deep learning. nature, 521, 436–444.

23. Rumelhart, D.E., Hinton, G.E. and Williams, R.J. (1986) Learning representations by back-propagating errors. nature, 323, 533–536.

24. Hochreiter, S. and Schmidhuber, J. (1997) Long short-term memory. Neural computation, 9, 1735–1780.

25. Graves, A. and Schmidhuber, J. (2005) Framewise phoneme classification with bidirectional LSTM and other neural network architectures. Neural networks, 18, 602–610.

26. Wang, Q. and Hao, Y. (2020) ALSTM: An attention-based long short-term memory framework for knowledge base reasoning. Neurocomputing, 399, 342–351.

27. Dey, R. and Salem, F.M. (2017), 2017 IEEE 60th international midwest symposium on circuits and systems (MWSCAS). IEEE, pp. 1597–1600.

28. Dos Santos, C. and Gatti, M. (2014), Proceedings of COLING 2014, the 25th International Conference on Computational Linguistics: Technical Papers, pp. 69–78.

29. Lai, S., Xu, L., Liu, K. and Zhao, J. (2015), Twenty-ninth AAAI conference on artificial intelligence.

30. Simonyan, K. and Zisserman, A. (2015) Very deep convolutional networks for large-scale image recognition. arXiv doi: https://arxiv.org/abs/1409.1556, 10 April 2015, preprint: not peer reviewed.

31. Wang, J., Yang, Y., Mao, J., Huang, Z., Huang, C. and Xu, W. (2016), Proceedings of the IEEE conference on computer vision and pattern recognition, pp. 2285–2294.

32. Vaswani, A., Shazeer, N., Parmar, N., Uszkoreit, J., Jones, L., Gomez, A.N., Kaiser, Ł. and Polosukhin, I. (2017), Advances in neural information processing systems, pp. 5998–6008.

33. Kitaev, N., Kaiser, Ł. and Levskaya, A. (2020), Proceedings of ICLR.

34. Choromanski, K., Likhosherstov, V., Dohan, D., Song, X., Gane, A., Sarlos, T., Hawkins, P., Davis, J., Mohiuddin, A. and Kaiser, L. (2022) Rethinking attention with performers. arXiv doi: https://arxiv.org/abs/2009.14794, 19 November 2022, preprint: not peer reviewed.

35. Wang, S., Li, B.Z., Khabsa, M., Fang, H. and Ma, H. (2020) Linformer: Self-attention with linear complexity. arXiv doi: https://arxiv.org/abs/2006.04768, 14 June 2020, preprint: not peer reviewed.

36. Roy, A., Saffar, M., Vaswani, A. and Grangier, D. (2021) Efficient content-based sparse attention with routing transformers. Transactions of the Association for Computational Linguistics, 9, 53–68.

37. Devlin, J., Chang, M.-W., Lee, K. and Toutanova, K. (2019), Proceedings of NAACL, pp. 4171–4186.

38. Safaya, A., Abdullatif, M. and Yuret, D. (2020), Proceedings of the Fourteenth Workshop on Semantic Evaluation, pp. 2054–2059.

39. Li, Y.-J., Zhang, H.-J., Pan, W.-M., Feng, R.-J. and Zhou, Z.-Y. (2021), Artificial Intelligence in China. Springer, pp. 524–530.

40. Nguyen, Q.T., Nguyen, T.L., Luong, N.H. and Ngo, Q.H. (2020), 2020 7th NAFOSTED Conference on Information and Computer Science (NICS). IEEE, pp. 302–307.

41. Huang, P., Zhu, H., Zheng, L. and Wang, Y. (2021), 2021 5th International Conference on Natural Language Processing and Information Retrieval (NLPIR), pp. 1–7.

42. Zhang, Z., Han, X., Liu, Z., Jiang, X., Sun, M. and Liu, Q. (2019), Proceedings of the 57th Annual Meeting of the Association for Computational Linguistics, pp. 1441–1451.

43. Kipf, T.N. and Welling, M.J.a.p.a. (2017), International conference on learning representations (ICLR ‘17).

44. Zhu, J., Cui, Y., Liu, Y., Sun, H., Li, X., Pelger, M., Yang, T., Zhang, L., Zhang, R. and Zhao, H. (2021), Proceedings of the Web Conference 2021, pp. 2848–2857.

45. Chen, J., Xie, Y., Wang, K., Wang, Z.H., Lahoti, G., Zhang, C., Vannan, M.A., Wang, B. and Qian, Z. (2018), International Conference on Medical Image Computing and Computer-Assisted Intervention. Springer, pp. 537–545.

46. Wang, K., Shen, W., Yang, Y., Quan, X. and Wang, R. (2020), Proceedings of the 58th Annual Meeting of the Association for Computational Linguistics, pp. 3229–3238.

47. Hamilton, W., Ying, Z. and Leskovec, J. (2017) Inductive representation learning on large graphs. Advances in neural information processing systems, 30.

48. Defferrard, M., Bresson, X. and Vandergheynst, P. (2016) Convolutional neural networks on graphs with fast localized spectral filtering. Advances in neural information processing systems, 29.

49. Wang, Z., Ye, X., Wang, C., Cui, J. and Philip, S.Y. (2020) Network embedding with completely-imbalanced labels. IEEE Transactions on Knowledge and Data Engineering, 33, 3634–3647.

50. He, X., Deng, K., Wang, X., Li, Y., Zhang, Y. and Wang, M. (2020), Proceedings of the 43rd International ACM SIGIR conference on research and development in Information Retrieval, pp. 639–648.

51. Brockschmidt, M. (2020), International Conference on Machine Learning. PMLR, pp. 1144–1152.

52. Ma, T., Dalca, A.V. and Sabuncu, M.R. (2022), Proceedings of the IEEE/CVF Winter Conference on Applications of Computer Vision, pp. 1933–1942.

53. Duan, W., He, X., Zhou, Z., Rao, H. and Thiele, L. (2021), Interspeech 2021, pp. 3216–3220.

54. Klicpera, J., Bojchevski, A. and Günnemann, S. (2019), 7th International Conference on Learning Representations.

55. Chen, J., Zhang, B., Xu, Y. and Wang, M. (2021) TextRGNN: Residual Graph Neural Networks for Text Classification. arXiv doi: https://arxiv.org/abs/2112.15060, 30 December 2021, preprint: not peer reviewed.

56. Wu, F., Souza, A., Zhang, T., Fifty, C., Yu, T. and Weinberger, K. (2019), International conference on machine learning. PMLR, pp. 6861–6871.

57. Lin, Y., Meng, Y., Sun, X., Han, Q., Kuang, K., Li, J. and Wu, F. (2021), Findings of the Association for Computational Linguistics: ACL-IJCNLP 2021, pp. 1456–1462.

58. Veličković, P., Cucurull, G., Casanova, A., Romero, A., Lio, P. and Bengio, Y. (2018), International conference on learning representations.

59. Wei, M., He, Y. and Zhang, Q. (2020), Proceedings of the 43rd International ACM SIGIR Conference on Research and Development in Information Retrieval, pp. 2367–2376.

60. Chandra, S., Mishra, P., Yannakoudakis, H., Nimishakavi, M., Saeidi, M. and Shutova, E. (2020) Graph-based modeling of online communities for fake news detection. arXiv doi: https://arxiv.org/abs/2008.06274, 23 November 2020, preprint: not peer reviewed.

61. Saito, T. and Rehmsmeier, M. (2015) The precision-recall plot is more informative than the ROC plot when evaluating binary classifiers on imbalanced datasets. PloS one, 10, e0118432.

62. Becht, E., McInnes, L., Healy, J., Dutertre, C.-A., Kwok, I.W., Ng, L.G., Ginhoux, F. and Newell, E.W. (2019) Dimensionality reduction for visualizing single-cell data using UMAP. Nature biotechnology, 37, 38–44.

63. Lundberg, S.M. and Lee, S.-I. (2017), Proceedings of the 31st international conference on neural information processing systems, pp. 4768–4777.

64. Richardson, B.C. (2002) Role of DNA methylation in the regulation of cell function: autoimmunity, aging and cancer. The Journal of nutrition, 132, 2401S–2405S.

65. Lv, H., Dao, F.-Y., Zhang, D., Guan, Z.-X., Yang, H., Su, W., Liu, M.-L., Ding, H., Chen, W. and Lin, H. (2020) iDNA-MS: an integrated computational tool for detecting DNA modification sites in multiple genomes. Iscience, 23, 100991.

66. Pan, X., Zuallaert, J., Wang, X., Shen, H.-B., Campos, E.P., Marushchak, D.O. and De Neve, W. (2020) ToxDL: deep learning using primary structure and domain embeddings for assessing protein toxicity. Bioinformatics, 36, 5159–5168.

67. Xu, H., Hu, R., Jia, P. and Zhao, Z. (2020) 6mA-Finder: a novel online tool for predicting DNA N6-methyladenine sites in genomes. Bioinformatics, 36, 3257–3259.

68. Chen, Z., Chen, Y.-Z., Wang, X.-F., Wang, C., Yan, R.-X. and Zhang, Z. (2011) Prediction of ubiquitination sites by using the composition of k-spaced amino acid pairs. PloS one, 6, e22930.

69. Chen, Z., Zhao, P., Li, F., Marquez-Lago, T.T., Leier, A., Revote, J., Zhu, Y., Powell, D.R., Akutsu, T. and Webb, G.I. (2020) iLearn: an integrated platform and meta-learner for feature engineering, machine-learning analysis and modeling of DNA, RNA and protein sequence data. Briefings in bioinformatics, 21, 1047–1057.

70. Liu, B., Liu, F., Wang, X., Chen, J., Fang, L. and Chou, K.-C. (2015) Pse-in-One: a web server for generating various modes of pseudo components of DNA, RNA, and protein sequences. Nucleic acids research, 43, W65–W71.

71. Liu, B., Liu, F., Fang, L., Wang, X. and Chou, K.-C. (2015) repDNA: a Python package to generate various modes of feature vectors for DNA sequences by incorporating user-defined physicochemical properties and sequence-order effects. Bioinformatics, 31, 1307–1309.

