## Supplementary Materials for "DeepBIO is an automated and interpretable deep-learning platform for biological sequence prediction, functional annotation, and visualization analysis"

### Supplementary Tables

**Supplementary Table 1. The detailed comparison between DeepBIO and other platforms.**

| Indicators | DeepBIO | BioSeq-BLM | iLearnPlus | BioSeq-Analysis2.0 | BioSeq-Analysis1.0 |
|---|---|---|---|---|---|
| Local-side applications or tools | × | √ | √ | √ | √ |
| Include traditional machine-learning models | × | √ | √ | √ | √ |
| Include deep-learning models | √ | √ | √ | × | × |
| GPU acceleration support | √ | √ | √ | × | × |
| Deal with imbalance datasets | √ | √ | × | √ | √ |
| Include pre-trained deep learning models | √ | × | × | × | × |
| Parameter optimization | √ | × | × | × | × |
| Multiple model comparison and visualization | √ | × | × | × | × |
| Biological sequence functional annotation | √ | × | × | × | × |
| The number of deep-learning methods | 42 | 6 | 7 | 0 | 0 |

**Supplementary Table 2. The construction of graph structures based on biological sequences.**

| Category Classic | Model |
|---|---|
| Graph-level | GCN |
| | GIN |
| | GAT |
| | GraphSage |
| | ChebGCN |
| | RECT-L |
| | LightGCN |
| | GNN-FILM |
| | HYPER-Conv |
| | HYPER-Attention |
| | APPNP |
| Node-level | TextGCN |
| | TextRGNN |
| | TextSGC |
| | BERTGCN |
| | BERTGAT |
| | RoBERTaGCN |
| | RoBERTaGAT |

We adopt two approaches for the graph structure construction based on the biological sequence including graph-level and node-level representations. For the graph-level representations, we transform the sequence information according to the smile format as a graph representation. Then we use several classical graph models to learn features. For the node-level representations, following the constructing method of TextGCN, we build a large and heterogeneous text graph which contains site-level elements (e.g., “ATCG” for DNA) as word nodes and the sequence as document nodes so that global word co-occurrence can be explicitly modeled and graph convolution can be easily adapted. The number of nodes in the text graph is the number of documents (corpus size) plus the number of unique words (vocabulary size) in a corpus. We simply set feature matrix as an identity matrix which means every word or document is represented as a one-hot vector as the input to graph model. We build edges among nodes based on word occurrence in documents (document-word edges) and word co-occurrence in the whole corpus (word-word edges). The weight of the edge between a document node and a word node is the term frequency-inverse document frequency (TF-IDF) of the word in the document, where term frequency is the number of times the word appears in the document, inverse document frequency is the logarithmically scaled inverse fraction of the number of documents that contain the word.

**Supplementary Table 3. The time cost of typical deep-learning models using DeepBIO on a training set that contains one thousand sequences.**

| Model-name | Time-cost(s) |
|---|---|
| DNN | 12.760 |
| GRU | 13.401 |
| RNN | 22.707 |
| LSTM | 14.356 |
| TextCNN | 597.950 |
| TextRCNN | 32.965 |
| VDCNN | 30.394 |
| GAT | 104.960 |
| GCN | 49.219 |
| Transformer | 26.379 |
| DNABERT | 801.366 |
| Reformer | 285.976 |
| Performer | 179.222 |
| Linformer | 257.919 |
| RoutingTransformer | 402.120 |

### Supplementary Figures

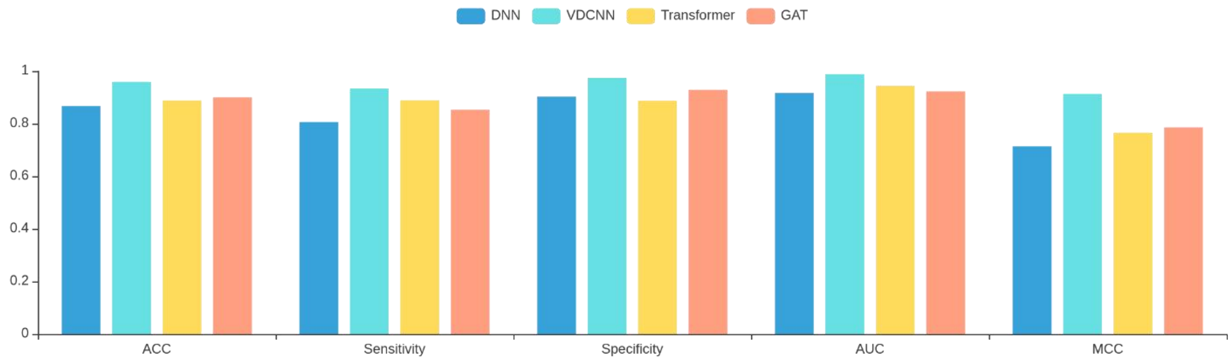


| Model Name | ACC | Sensitivity | Specificity | AUC | MCC |
|---|---|---|---|---|---|
| DNN | 0.866 | 0.806 | 0.902 | 0.917 | 0.713 |
| VDCNN | 0.959 | 0.934 | 0.974 | 0.987 | 0.913 |
| Transformer | 0.887 | 0.888 | 0.887 | 0.944 | 0.765 |
| GAT | 0.900 | 0.853 | 0.928 | 0.922 | 0.786 |

**Supplementary Figure 1. Performance of deep learning models on the prediction of protein toxicity.**

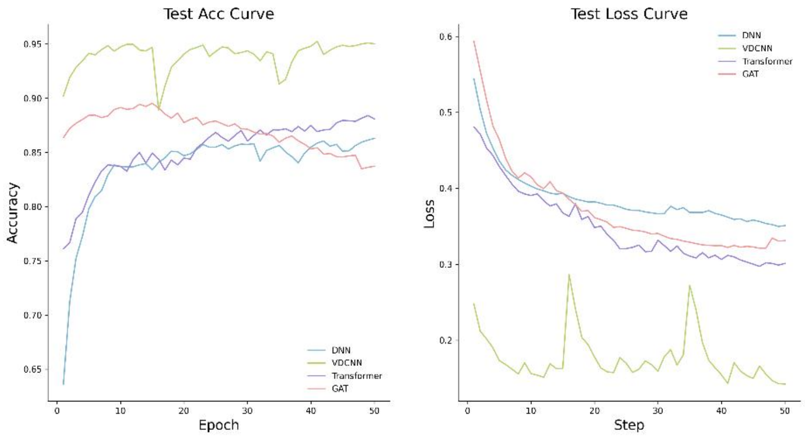


**Supplementary Figure 2. Epoch plot on the prediction of protein toxicity.**

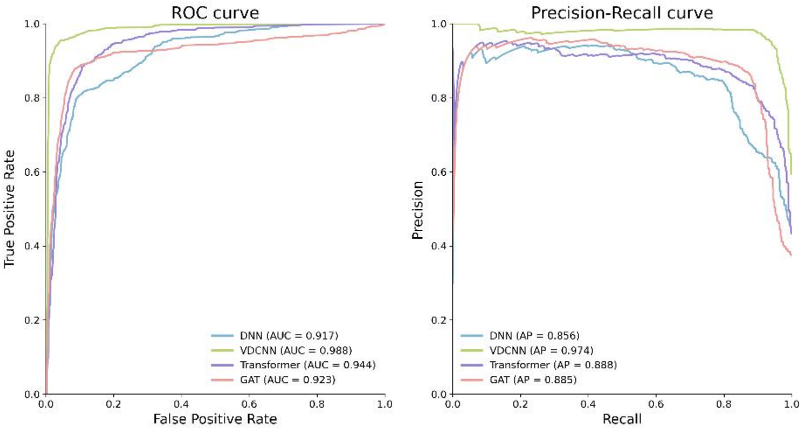


**Supplementary Figure 3. ROC and PR curve on the prediction of protein toxicity.**

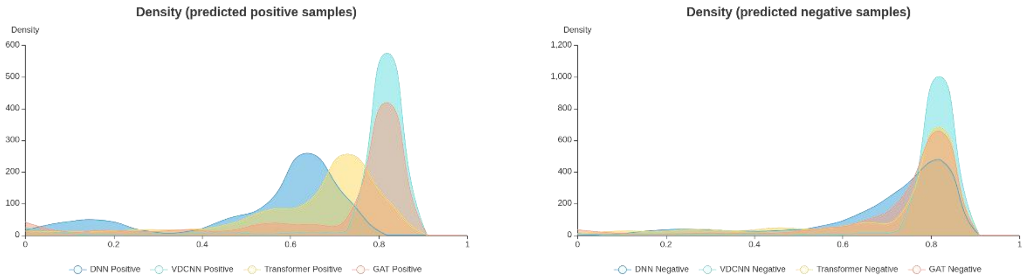


**Supplementary Figure 4. Density distribution of the prediction confidence by different deep learning models on the prediction of protein toxicity.**

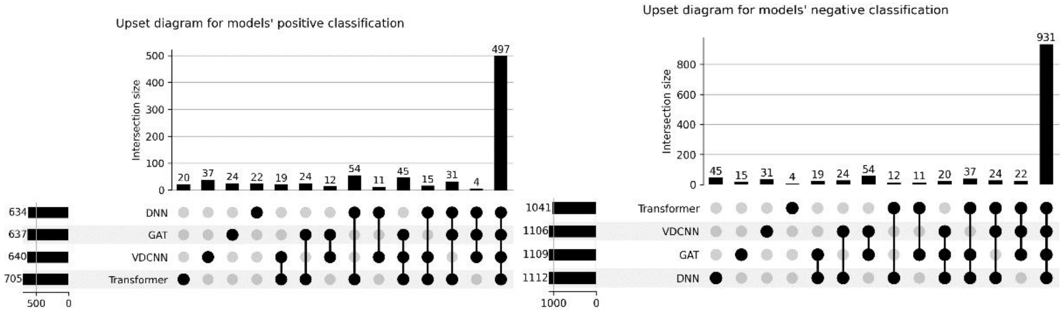


**Supplementary Figure 5. Upset diagram on the prediction of protein toxicity.**

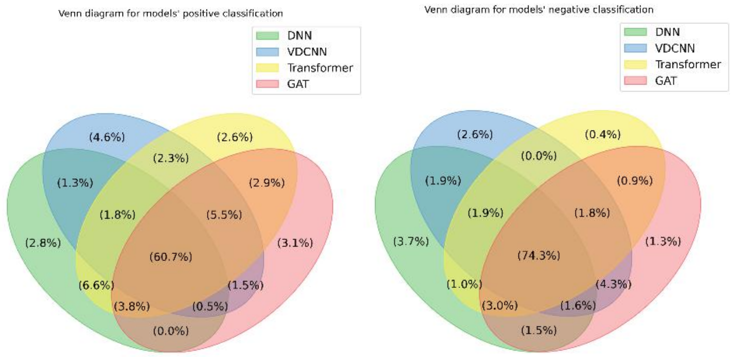


**Supplementary Figure 6. Venn diagram on the prediction of protein toxicity.**

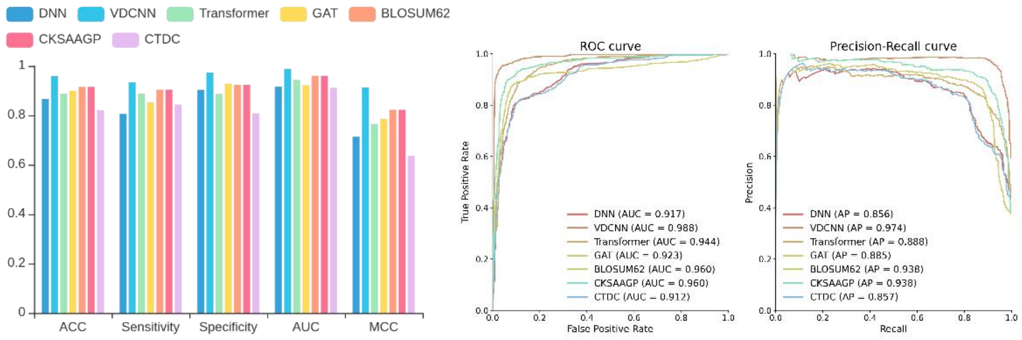


**Supplementary Figure 7. Feature performance comparison with hand-crafted features on the prediction of protein toxicity.**

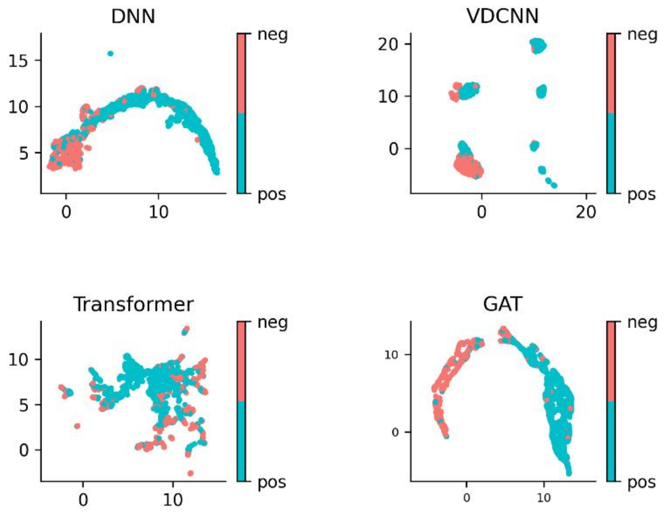


**Supplementary Figure 8. Feature space visualization by UMAP on the prediction of protein toxicity.**

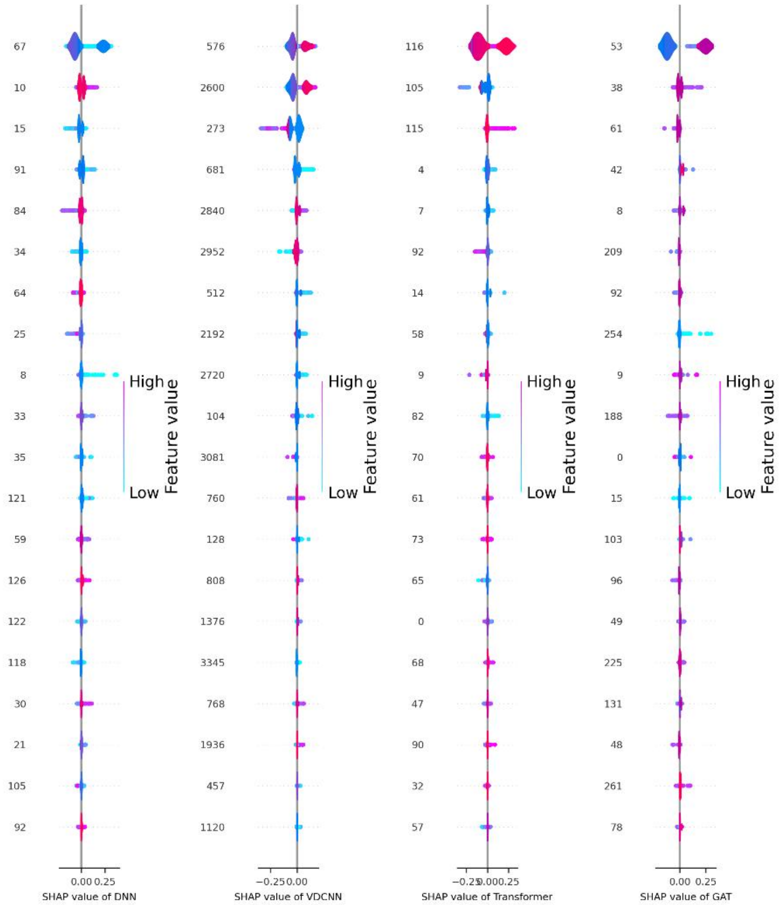


**Supplementary Figure 9. Feature importance analysis by SHAP on the prediction of protein toxicity.**

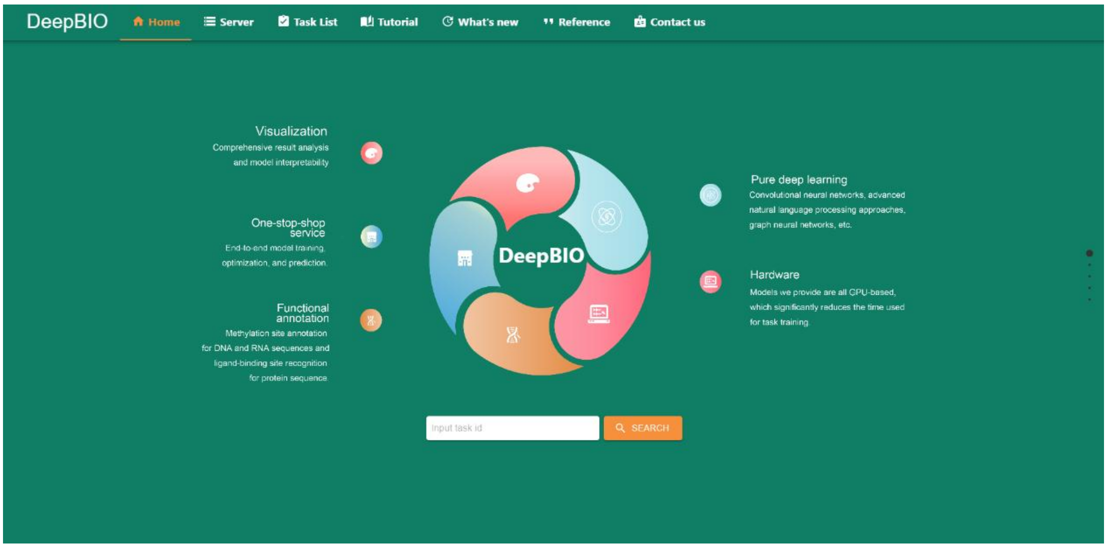


**Supplementary Figure 10. Home page of our DeepBIO online server.** It shows the overall functions of our DeepBIO platform.

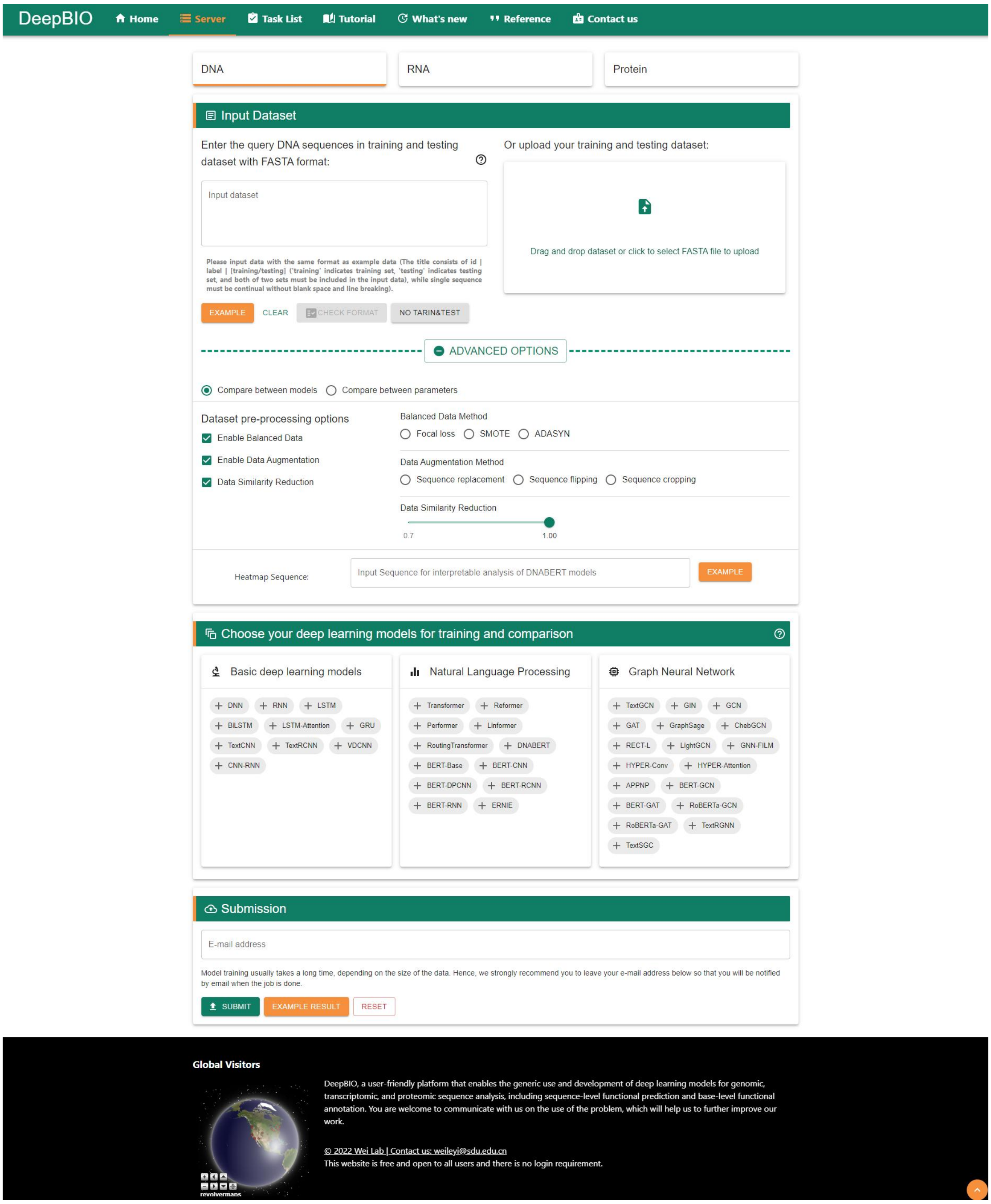


**Supplementary Figure 11. Deep learning based prediction module of our DeepBIO online server.** Users can train and evaluate deep learning models based on their own biological sequence data.

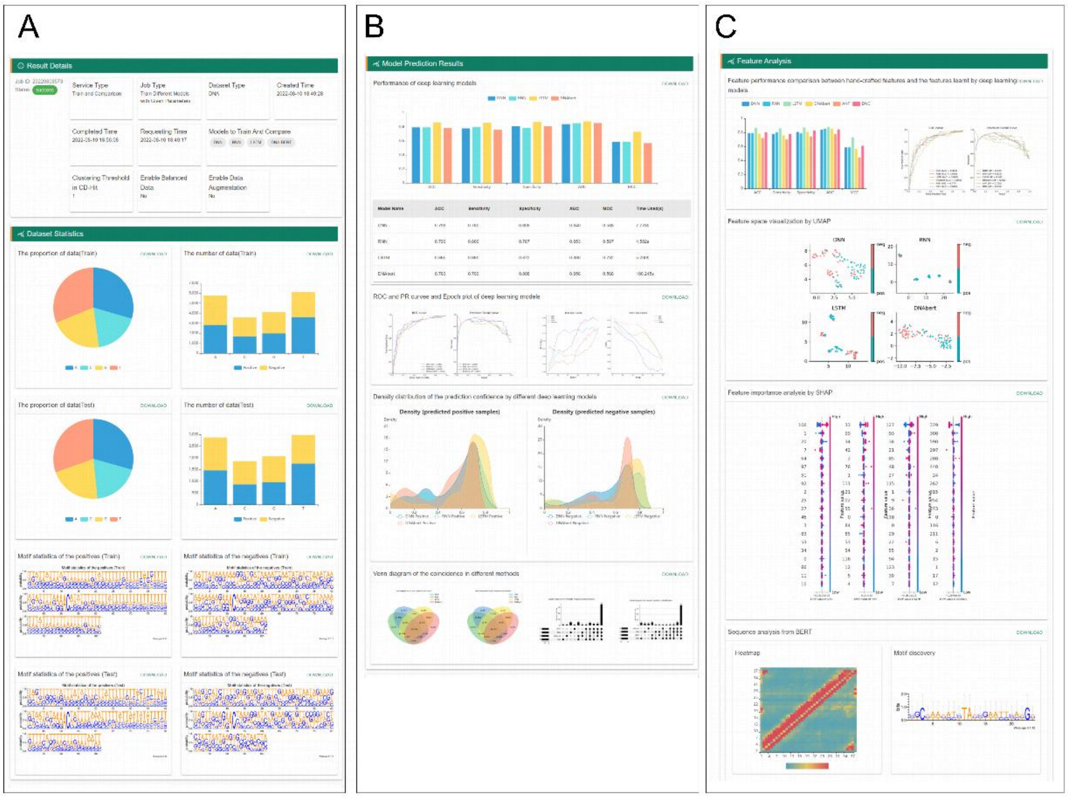


**Supplementary Figure 12. Result report of deep learning based prediction module using DeepBIO.** Here, we choose the DNA example data on our online server as case datasets to show the comprehensive visualization and analysis results.

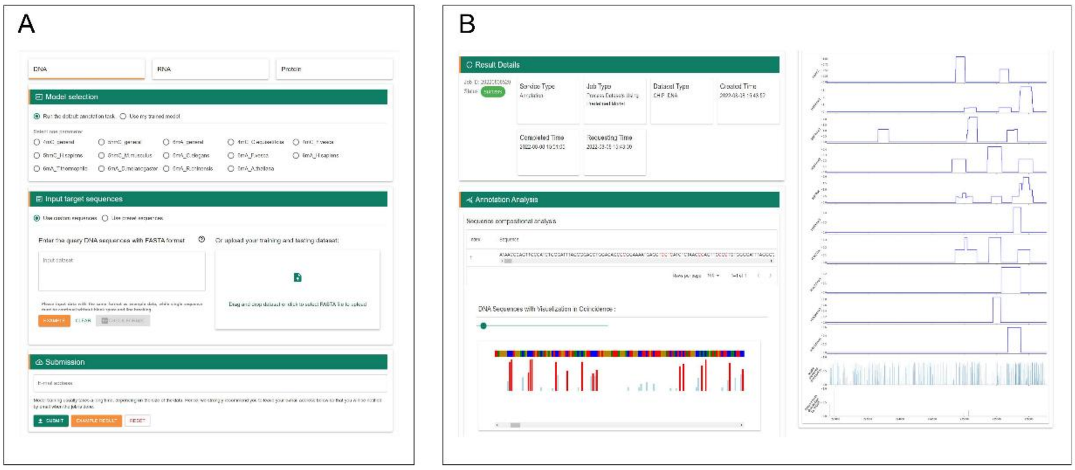


**Supplementary Figure 13. (A) Sequence functional annotation module of our DeepBIO online server for DNA.** Users can select our default models or the models they have trained in previous tasks for biological sequence functional annotation. **(B) Result report of sequence functional annotation module using DeepBIO for DNA methylation site prediction.** Here, we select a segment of our preset human gene by choosing a specific cell line and chromosome to show the comprehensive visualization analysis between DNA methylation annotation results and different histone modification signals.

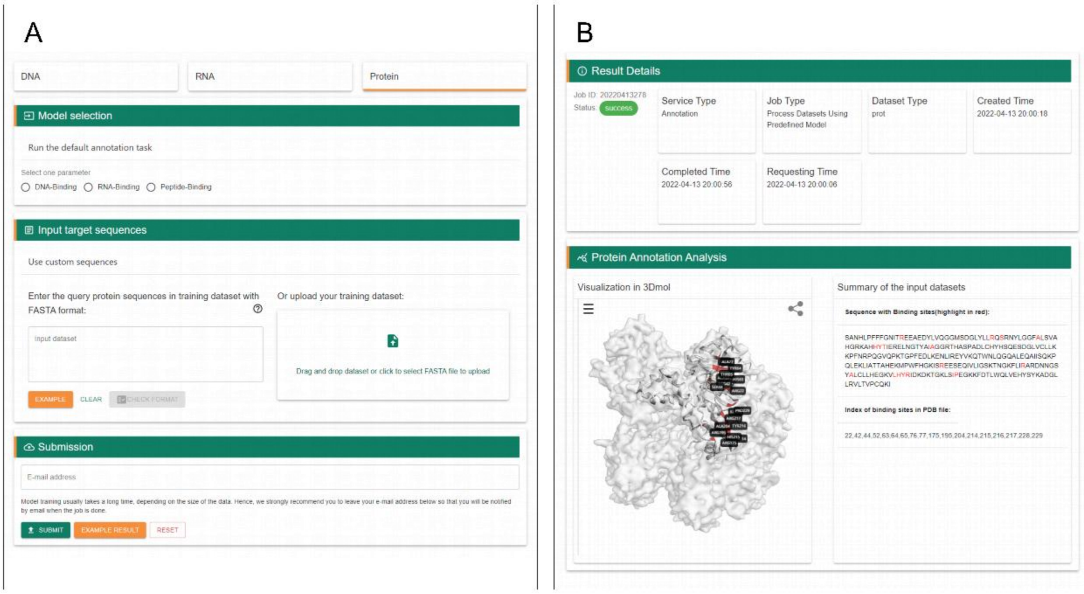


**Supplementary Figure 14. (A) Sequence functional annotation module of our DeepBIO online server for protein.** Users can choose one of three different tasks (DNA-binding prediction, RNA-binding prediction, and Peptide-binding prediction) for protein-binding site annotation. **(B) Result report of sequence functional annotation module using DeepBIO for protein-peptide binding site annotation.** Here, to intuitively show the protein-peptide binding site annotation results, we visualize and annotate the binding residues on the 3D structure of a given protein sequence with PDB ID.

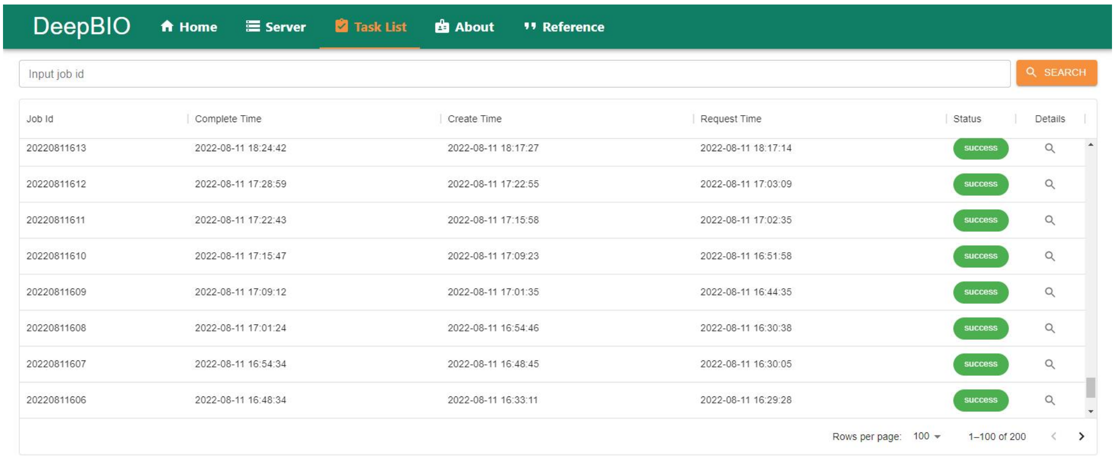


**Supplementary Figure 15. Task list page of our DeepBIO online server.** Users can obtain their result report according to the relevant job id.
